# Understanding Chicks’ Emotions: Are Eye Blinks & Facial Temperatures Reliable Indicators?

**DOI:** 10.1101/2022.01.31.478468

**Authors:** Naomi Pijpers, Huib van den Heuvel, Ian H. Duncan, Jessica Yorzinski, Suresh Neethirajan

## Abstract

In commercial farming systems, chicks are reared without a mother. This absence of maternal influence can cause welfare problems when the chicks become older. Chicks imprint on their mothers they are young, and this mediates their stress and fear response. It is important to recognise problems early in the development of chicks to avoid welfare issues when they are older. One way to assess welfare is by measuring affective states. Research has shown chickens can display empathy, both towards their offspring and towards conspecifics. Measures of negative and positive affective states, either behavioural or physiological, could be good welfare indicators. This study employed non-invasive methods to measure affective states in laying hen chicks. Using video and thermal imaging, it analysed temperature changes in the peripheral areas and head region as well as changes in blinking behaviour before and after exposure to a stressor. The prediction was that the temperature would decrease in the eye and peripheral regions in response to a stressor and that the blinking rate would decrease. These changes would be indicative of a negative affective state. The results showed that the eye temperature as well as the blinking rate both decreased, whereas the temperature in the head region and the beak area increased. These results could be indicative of a negative affective state.

## 1 Introduction

In commercial farming systems, laying hen chicks are not brooded by a mother hen but in large incubators in hatcheries and brooders on the farm (1). This means the chicks will never have a mother figure and will not benefit from maternal care (2). Chicks are precocial animals, meaning they are mobile from the moment they are hatched, but they still require maternal care in the first weeks of their lives (3). This maternal care positively influences the behavioural development of the chicks, for example, by imprinting certain behaviour related to vocalisations, feeding, and mediating fear and stress responses (4). How the chicks are reared from hatching until laying age can influence the welfare of the grown laying hens. Laying hen chicks are exposed to many stress factors in early life, such as transportation, vaccination, manual sex sorting, not to mention the hatching in the incubators itself (5). Earlier studies have found that mother hens can buffer the stress of their chicks and that chicks show less fearful responses in the presence of their mother (2). A bird in a good state of welfare should display a healthy body and a positive affective state and should be able to express natural behaviour (6). Early negative experiences can have long-term effects on the development and behaviour of the laying hens and can cause unwanted behaviour, like feather pecking (7).

However, natural brooding is not commercially feasible in domestic chicken farms (8). Brooding hens have a lower feed conversion (9) and their growth rate is reduced (10). In addition, brooding hens do not lay eggs, therefore, taking up space that could be used for hens that are producing eggs (4). Thus, it is important to find other methods to fulfil the needs of the chicks that their mother would provide for naturally and to improve the chicks’ welfare. To develop a welfare platform with the help of sensors and technology, many aspects can be researched, one of which is the assessment of affective states. A deeper understanding of affective states in animals is needed to take the proper measures required for positive states of welfare and to limit negative states welfare (11).

In animals, affective states can be measured through physiological components and behavioural measurements, like skin temperature, head movements, and locomotion (12,13). Affective states can be divided into arousal, the intensity of the emotion and valence, and whether the emotion is negative or positive. In avian species, both affective states of positive and negative valence have been observed. Positive affective states, such as the feeling of pleasure, are important for animal welfare (14). Negative affective states, such as fear and anger, can be useful for survival, for example, when dangerous situations arise and can also be an indicator of welfare (15).

A study on the presence of positive affective states in Japanese quail found that changes in several facial cues (throat feather angle, pupil dilation, and crown feather height) in response to rewarding behaviour, could be an indicator of positive emotions (16). In an experiment with broiler chickens, the effects of environmental enrichment were investigated. The presence of positive affective states was investigated with the help of a judgement bias test. It was found that more complex environments contributed to positive affective states in the chickens (17). Anderson et al., (18) also studied the anxiety, the reaction to a perceived threat, and fear, the reaction to a known threat, of broiler chickens in response to stocking density (19). It was found that high stocking densities contributed to a reduction of the fear response and complex environment reduced anxiety. An overview of affective states measured in avian animals with sensors can be found in Table 1.

Emotions are not individual components. In many social animals, the phenomenon of emotional contagion has been found. Emotional contagion can be defined as the emotional state-matching of a subject with an object (20). It means the individual is aware of the emotions of a conspecific animal (11). In chickens, it was measured that the stress response of an observer chicken was greater when the demonstrator chicken was handled roughly than when it was handled gently, suggesting the presence of emotional contagion and empathy (21). Chickens do not have to experience a stressful situation to feel stressed. The mere presence of a stressed conspecific can cause them to feel stressed (11).

Important for emotional contagion is socially mediated arousal, which is the increased alertness to respond that can occur when one animal perceives another’s behaviour or physiology (22). Behavioural indicators can be freezing responses and reduced maintenance behaviour, such as preening. Physiological indicators can be heart rate variability, temperature variations, and hormonal variations. A study about the presence of socially mediated arousal in mother hens, by Edgar et al. (2015) (2), found that when chicks received a mild stress treatment, the mothers had an increased heart rate, experienced stress-induced hypothermia, produced more vocalisations, increased standing alert, and reduced preening behaviour.

Although this study does not focus on emotional contagion, but rather on specific affective states, it is important to mention that chickens can feel for one another, which has implications for the welfare of the chickens.

**Table 1:**
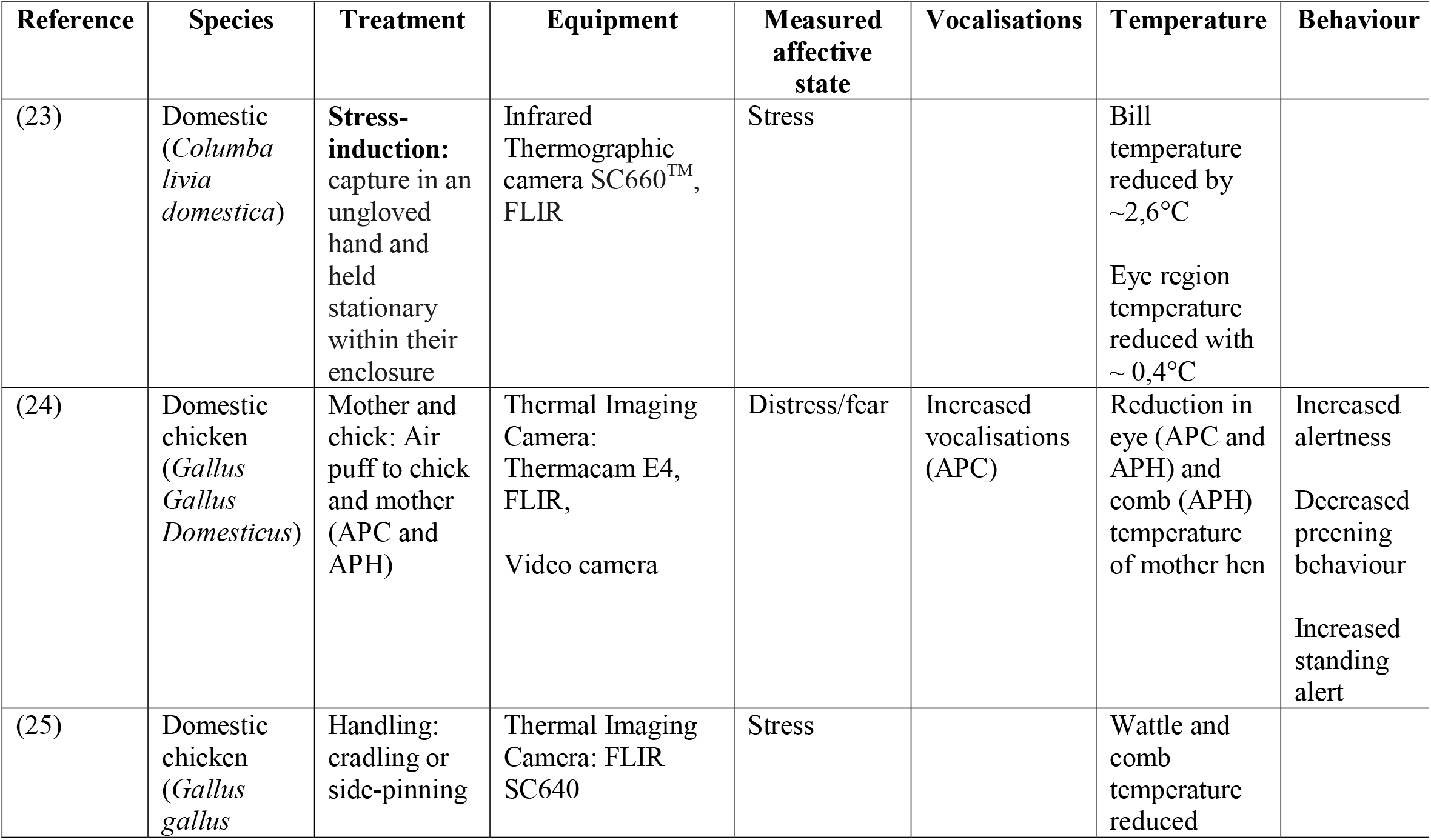
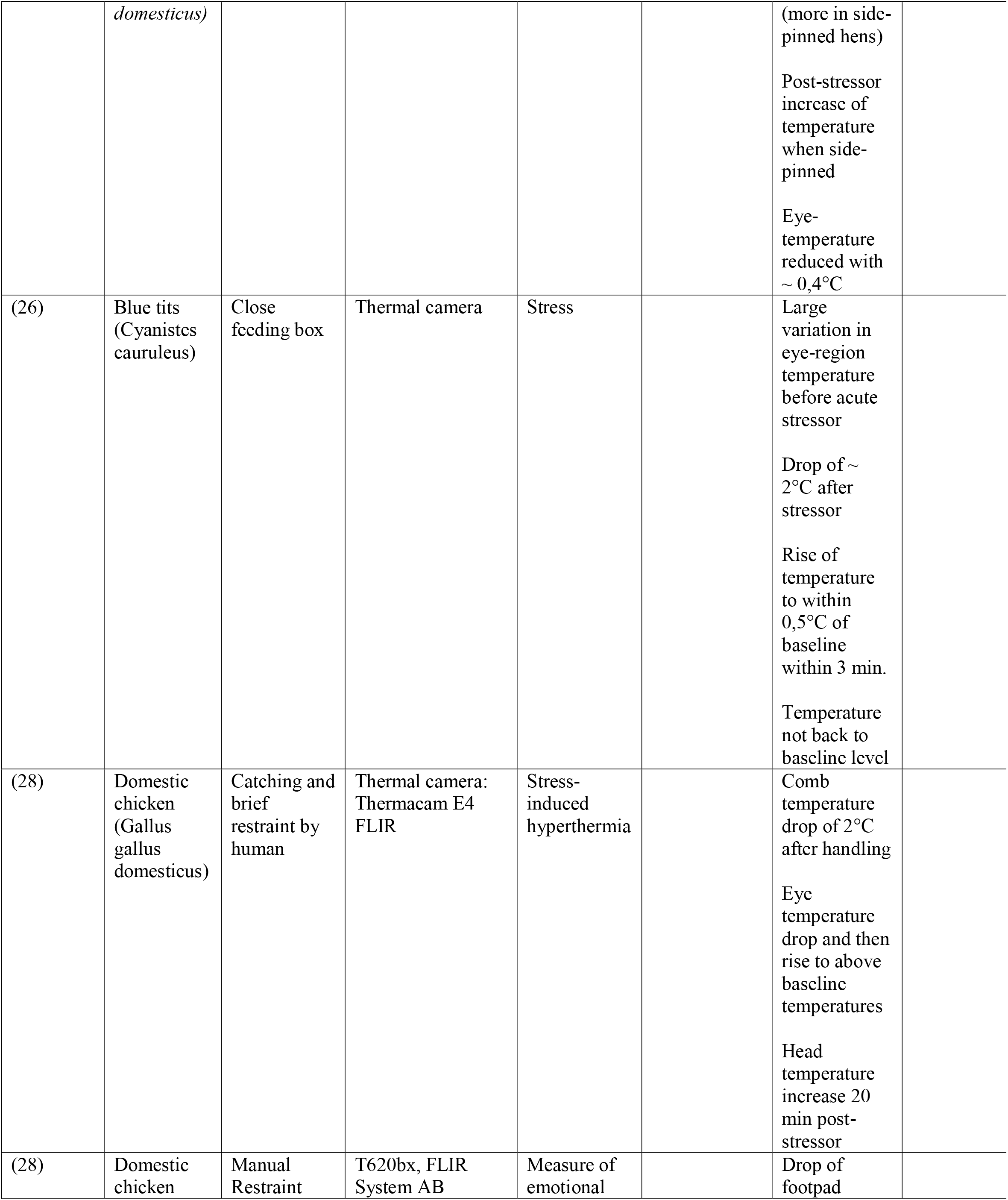
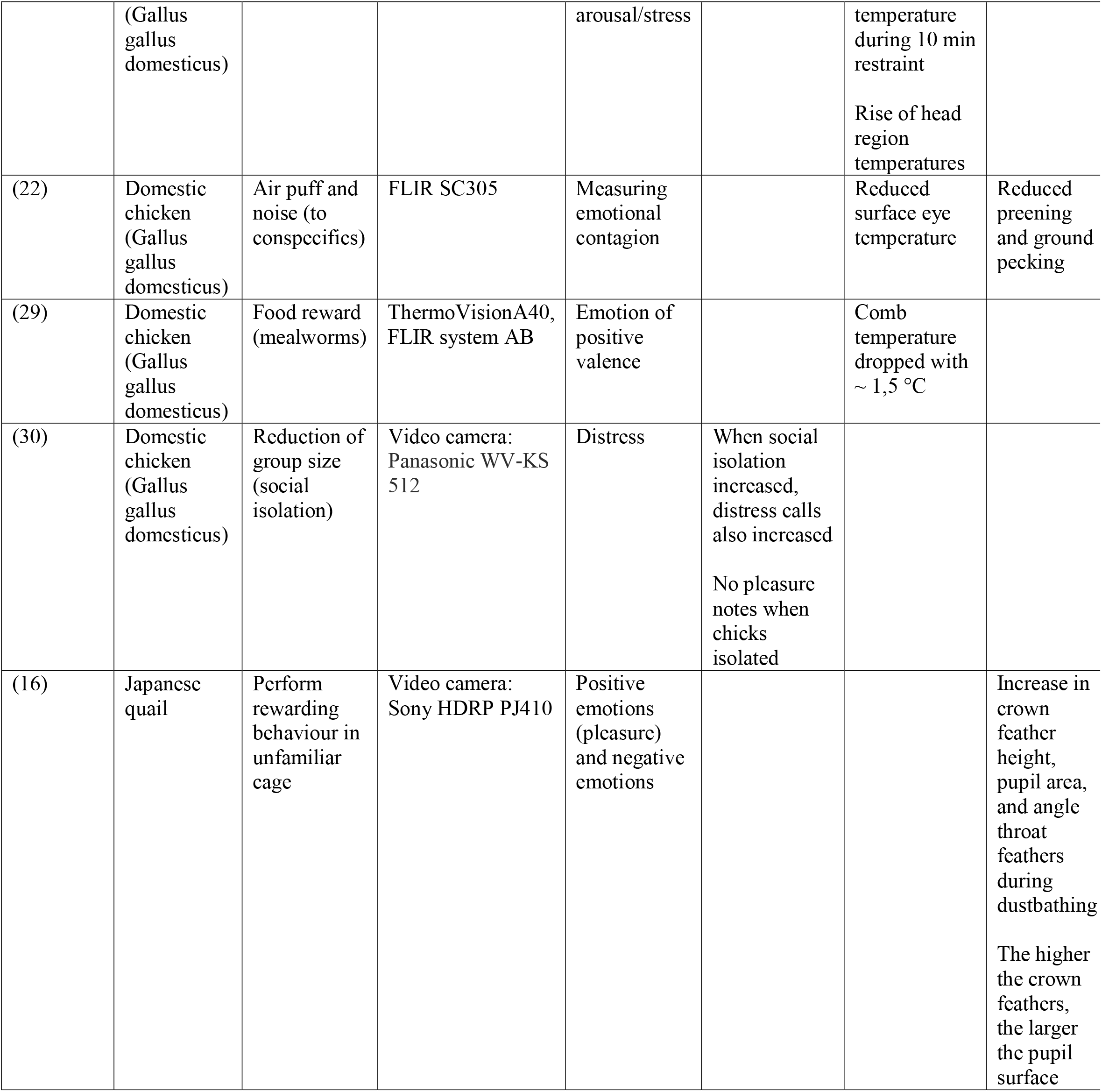
Affective states measured in birds with sensors and imaging technologies

| Reference | Species | Treatment | Equipment | Measured affective state | Vocalisations | Temperature | Behaviour |
| --- | --- | --- | --- | --- | --- | --- | --- |
| (23) | Domestic (*Columba livia domestica*) | **Stress-induction:** capture in an ungloved hand and held stationary within their enclosure | Infrared Thermographic camera SC660™, FLIR | Stress |  | Bill temperature reduced by ~2,6°C<br>Eye region temperature reduced with ~ 0,4°C |  |
| (24) | Domestic chicken (*Gallus Gallus Domesticus*) | Mother and chick: Air puff to chick and mother (APC and APH) | Thermal Imaging Camera: Thermacam E4, FLIR,<br>Video camera | Distress/fear | Increased vocalisations (APC) | Reduction in eye (APC and APH) and comb (APH) temperature of mother hen | Increased alertness<br>Decreased preening behaviour<br>Increased standing alert |
| (25) | Domestic chicken (*Gallus gallus*) | Handling: cradling or side-pinning | Thermal Imaging Camera: FLIR SC640 | Stress |  | Wattle and comb temperature reduced |  |
| Non-Invasive Indicators of Stress in Chickens |  |  |  |  |  |  |  |
|  | *domesticus*) |  |  |  |  | (more in side-pinned hens)<br>Post-stressor increase of temperature when side-pinned<br>Eye-temperature reduced with ~ 0,4°C |  |
| (26) | Blue tits (Cyanistes caeruleus) | Close feeding box | Thermal camera | Stress |  | Large variation in eye-region temperature before acute stressor<br>Drop of ~ 2°C after stressor<br>Rise of temperature to within 0,5°C of baseline within 3 min.<br>Temperature not back to baseline level |  |
| (28) | Domestic chicken (Gallus gallus domesticus) | Catching and brief restraint by human | Thermal camera: Thermacam E4 FLIR | Stress-induced hyperthermia |  | Comb temperature drop of 2°C after handling<br>Eye temperature drop and then rise to above baseline temperatures<br>Head temperature increase 20 min post-stressor |  |
| (28) | Domestic chicken | Manual Restraint | T620bx, FLIR System AB | Measure of emotional |  | Drop of footpad |  |
|  | (Gallus gallus domesticus) |  |  | arousal/stress |  | temperature during 10 min restraint<br>Rise of head region temperatures |  |
| (22) | Domestic chicken (Gallus gallus domesticus) | Air puff and noise (to conspecifics) | FLIR SC305 | Measuring emotional contagion |  | Reduced surface eye temperature | Reduced preening and ground pecking |
| (29) | Domestic chicken (Gallus gallus domesticus) | Food reward (mealworms) | ThermoVisionA40, FLIR system AB | Emotion of positive valence |  | Comb temperature dropped with ~ 1,5 °C |  |
| (30) | Domestic chicken (Gallus gallus domesticus) | Reduction of group size (social isolation) | Video camera: Panasonic WV-KS 512 | Distress | When social isolation increased, distress calls also increased<br>No pleasure notes when chicks isolated |  |  |
| (16) | Japanese quail | Perform rewarding behaviour in unfamiliar cage | Video camera: Sony HDRP PJ410 | Positive emotions (pleasure) and negative emotions |  |  | Increase in crown feather height, pupil area, and angle throat feathers during dustbathing<br>The higher the crown feathers, the larger the pupil surface |

### 1.1 Blinking behaviour

Blinking rate as a measure of emotions has been tested in several animal species. In horses, the spontaneous blinking rate was measured as a possible indicator of stress. It was found that the blinking rate first decreased and after a few minutes increased, a result similarly found in humans (31). A study with crows showed that when crows are exposed to a threat, their blinking rate decreases, as they respond with a fixed gaze (32). Dogs showed an increased blinking rate in response to fear and frustration, although this might have been a response to stress instead of these specific emotions (33).

Blinking is necessary for clear vision, but it also temporarily impairs vision, as visual information input is inhibited (34,35). When an animal blinks and closes its eye, it is essentially blind and can not pay attention to its surroundings. A high blinking rate and long duration of blinks can cause intermittent blindness, which leads to a reduced detection of threats (36). Chickens have mostly monocular vision, with the field of view being around 300°, with just 30° overlap, thus allowing for binocular vision. Gaze shifts are needed to obtain large changes in view, which are enabled by the flexible neck and light head (37). A study tested whether songbirds were able to modify their blinking behaviour when they were subjected to a potential threat. It was found that the birds modified their blinking behaviour based on reactivity, and they could modify their blinking rate based on a perceived risk (34).

Measured with the help of video cameras, the blinking behaviour could also be an interesting indicator to explore affective states in chicks. When an animal is in a negative affective state, for example, when it is frightened, its blinking behaviour might change. Due to the increased alertness, the blinking rate might decrease. Blinking could therefore be a measure of affective state.

### 1.2 Thermal imaging

Another way to assess affective states in chickens is by measuring physiological responses, such as temperature (38). Infrared thermography is used as a non-invasive technique to measure body surface temperature which in turn can relate to stress and emotions in hens (28). Physiological components can give indicators of emotional states that cannot be assessed verbally in animals. An animal under stress will display cutaneous vasoconstriction, leading to a drop in skin temperature. This is paired with an increase in core temperature, which in turn is followed by vasodilation, resulting in a post-stressor rise in peripheral temperature. This is known as emotional fever (28).

It has been shown that shank temperature of laying hens drops by 1-2°C for one to two minutes when they are exposed to a frightening visual stimulus (39).

Infrared thermography, or thermal imaging, can detect infrared radiation emitted from an object (40). It is used in many animal studies related to stress, emotional arousal, and animal welfare (15). To measure skin temperature with infrared thermography, the skin needs to be bare. In chickens, the wattles, comb, and head are already bare, which makes the chicken a good model for comparing temperatures (25).

A study showed that when chickens were handled, stress caused an initial surface comb and eye temperature drop of 2 °C and 0.8 °C respectively after one minute of handling, while core temperature increased (41). Research in aggressive behaviour in pheasants found the head temperature of both the attacker and the recipient of the attack first decreased before the attack, and afterward showed an increase (42). Another study found that a rougher handling procedure also caused a larger temperature shift, which suggests that skin temperature may be a way of quantifying by measures of stressor intensity (25).

### 1.3 Hypothesis

In this study we examine whether a negative affective state can be observed in laying hen chicks in response to a stressor by measuring physiological and behavioural components with the help of video and thermal imaging. It is predicted the eye temperature would drop, whereas the overall head temperature will rise as a sign of distress or fear. It is also predicted that the blinking behaviour will change in response to a stressor, such as a reduced blinking rate to increase alertness.

## 2 Materials and methods

We examined the blinking behaviour and temperature changes in laying hen chicks as a measure of affective states during a non-invasive experiment. The chicks were investigated for two weeks, between November and December 2021, at the Research Facility CARUS of the Wageningen University & Research.

### 2.1 Subjects

Fifty laying hen chicks of the breed Super Nick were used for this experiment. Upon arrival, they were one day old. The experiment started when they were nine days old. The chicks were housed in one pen of 2×4 meters (n=10) and two 4×4 meter pens (n=20). The chicks were only handled when they were brought in, but not during the experiment. They received the necessary vaccinations upon arrival.

### 2.2 Procedure

We assessed the chicks two or three times a week for two weeks. Their exact age on the days of assessment was respectively: 9 days, 11 days, 15 days, 17 days, and 19 days. The experiment had within-treatment control. The chicks were assessed before inducing a stressor and during or after inducing a stressor. The assessment was done by taking thermal pictures and taking close-up videos to measure the blinking behaviour. The stressors used were a playback of sounds and a visual stressor. For the thermal pictures, only the visual stressor was used. The visual stressor was a pink umbrella opening close to the pen. The umbrella was only opened and closed once per cage after which thermal pictures were immediately taken. On days 9, 11, 15, 17, and 19 the chicks received the umbrella stressor. During the assessment of the blinking behaviour, the chicks received two different treatments. On days 9, 11, and 15, the chicks received only the visual stressor, the opening and closing of the pink umbrella. On days 17 and 19, the chicks received only the sound stressor which was a playback of the aggressive barking of a dog. This sound was played on a speaker (JBL Flip 2). For this stressor, the chicks were filmed while the sound was playing. This sound was played for 10 minutes per cage during which the birds were filmed. It was measured whether these stressors evoked a certain response, both physiological and behavioural.

For the thermal pictures, a thermographic camera was used, the FLIR 1020. The pre-stress thermal pictures of all chicks were taken for 45 minutes. Fifteen pictures were taken in the cages where 20 birds were housed, and 10 pictures were taken in the cage where 10 birds were housed. Then stress was induced. Immediately afterward, 40 thermal pictures of all the chicks were taken again for 45 minutes. Then stress was induced again. One hour after taking pictures after the stress treatment, 40 post-stress thermal pictures were taken. In total, 40 pictures were taken in each condition: pre-stress, stress, and post-stress, 120 pictures per day.

In the experiment, we planned to measure the temperature of the eye, comb, head, and beak area, so parts of the chicken are bare-skinned. However, the chicks did not have a comb yet, so we could only measure the eye, beak, and head temperature. The thermal camera was handled manually. The head of the chicks was photographed, and the photos were manually analysed with the program FLIR ResearchIR. In the program, the region of interest could be drawn manually and provided the average and maximum temperature of the area (Figure 1). The ambient temperature of the room the chicks were housed in decreased as the chicks got older. The maximum temperatures on the different days of assessment were: 31.2 °C on day 9, 30.9 °C on day 11, 30.2°C on day 15, 30.3 °C on day 17, and 27.9 °C on day 19 in the cages with 20 chicks. The ambient temperatures of the cage where 10 chicks were housed were: 31.0 °C on day 9, 39.8 °C on day 11, 31.3 °C on day 15, 31.3 °C on day 17, and 27.9 °C on day 19. Statistical calculations were performed in SPSS.

**Figure 1:**
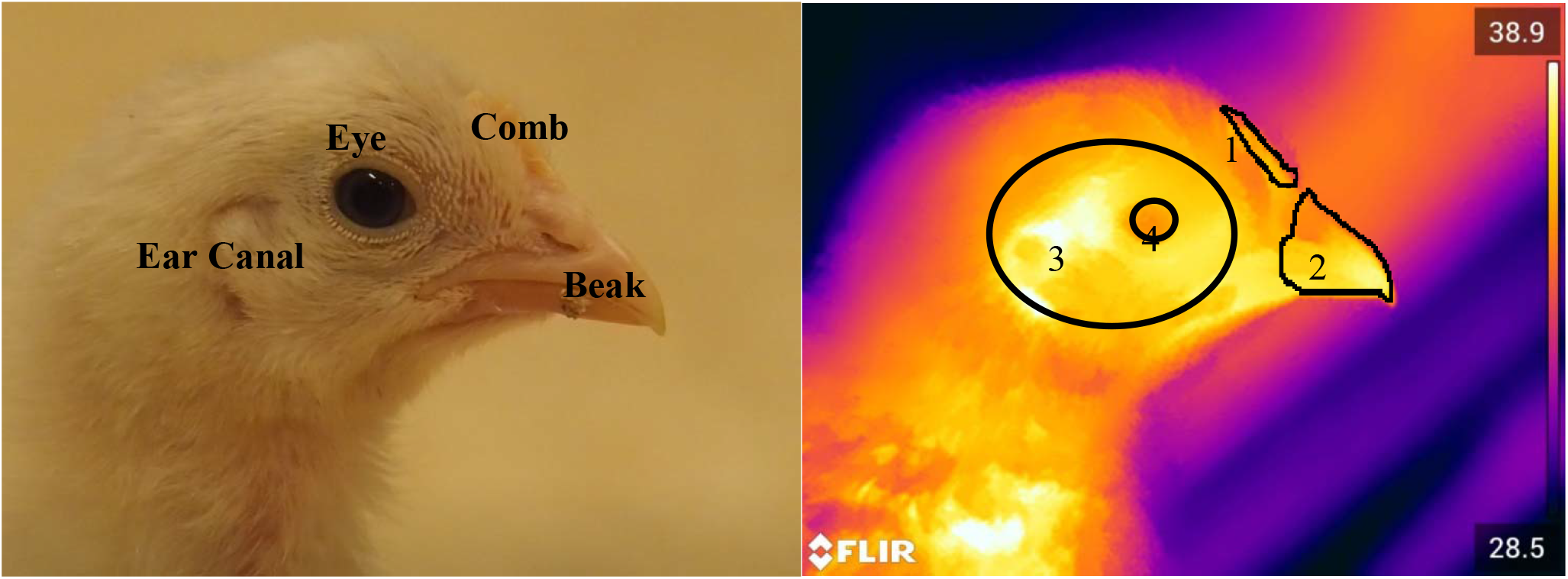
An example of the studied chick. Regular picture (left) and thermal picture (right), with the regions of interest (ROI) encircled. 1: comb, 2: beak, 3: head, 4: eye.

The blinking behaviour of the laying hen chicks was assessed as well. This was done with a video camera (Olympus OM-D E-M10 Mark II, 50 fps, 16 megapixels resolution), placed on a tripod for stability. In addition, a GoPro Hero 9 camera with the frame rate of 240 fps was also deployed to collect images of the chicks. This camera was used to capture more elaborate and in-depth details on the face of the chicks. The blinking behaviour was measured before and after exposure to the stressor. As the experiment was non-invasive, picking up the birds was not allowed, and, therefore, it was not possible to assess the blinking behaviour of each chick from a particular position and distance. Instead, around five birds in each pen were followed, and their eyes were filmed as closely as possible for 1 to 3 minutes. The birds did not wear an identification tag, so the observer was not able to identify each bird. However, the research ensured that the same bird was not filmed twice by filming each chick in a different area of the cage. In addition, some birds could be identified by an external component such as the presence of faeces on the wings or the rectum. The eye closures were divided into two categories: (1) The partial closure of the upper and under the eyelid and the full closure of the nictitating membrane (partial blink; PB) and (2) the full closure of the upper and under the eyelid (full blink; FB) (Figure 3).

**Figure 2:**
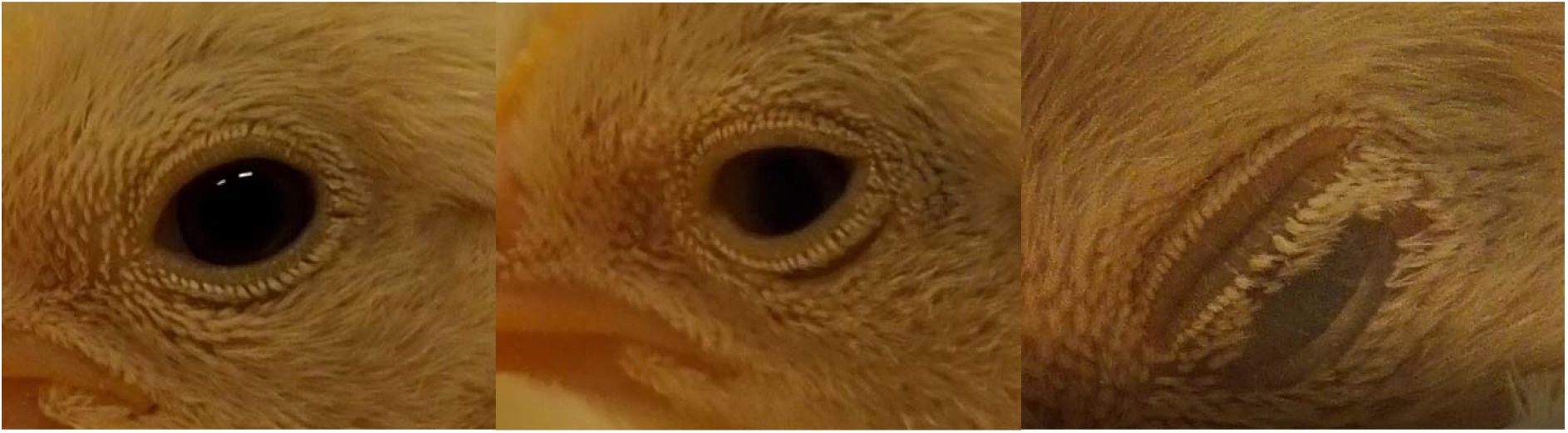
Overview of the types of chick blinks. From left to right: open eye, partial blink (PB), and full blink (FB)

**Figure 3:**
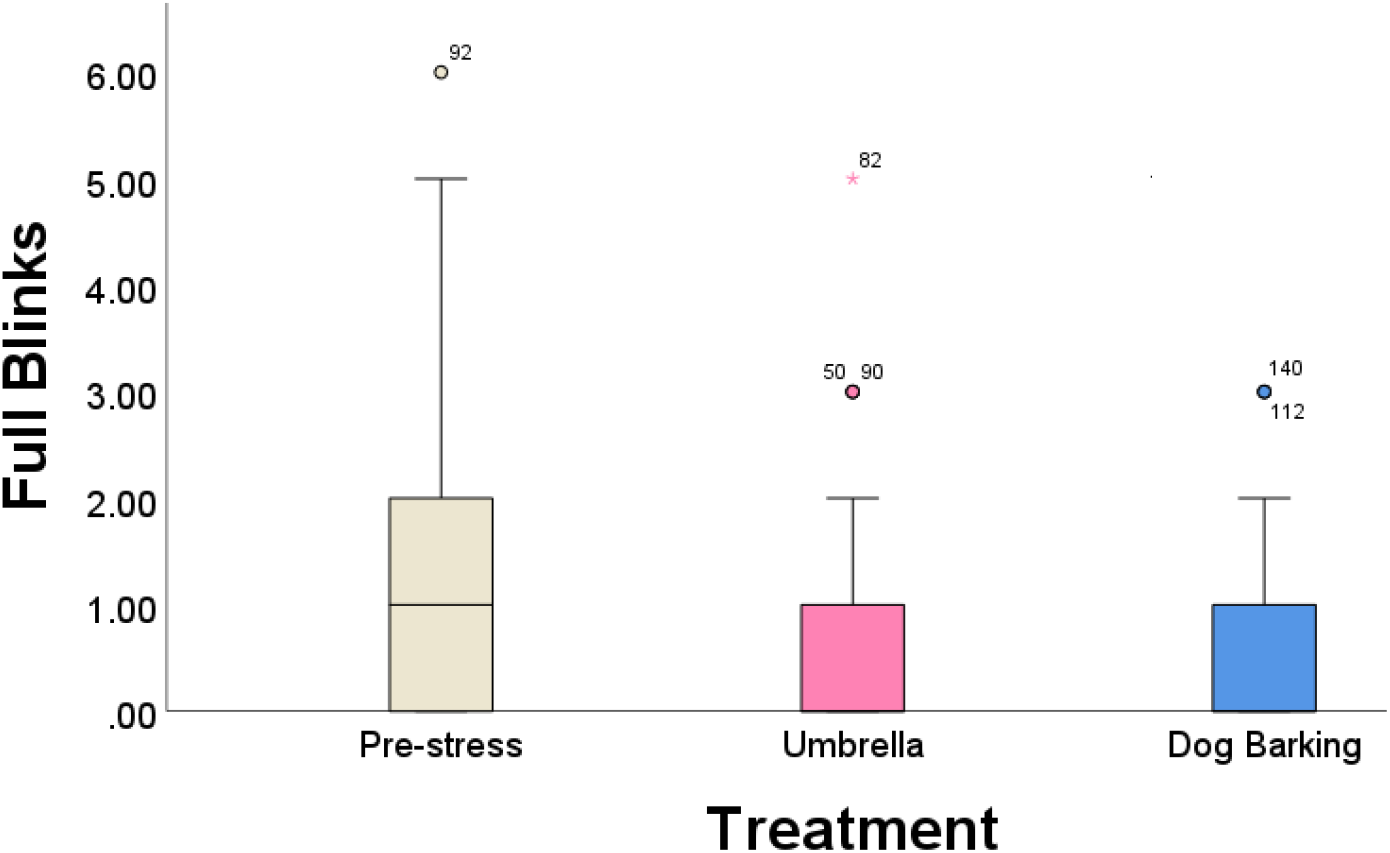
The number of full blinks during the different treatments (Umbrella and Dog Barking). Before exposure to the stressor, the chicks displayed a higher number of full blinks, and, thus, they blink less when they are stressed (p = 0.0211). Full blinks were twice more present when the birds were non-stressed than after the two different treatments of the umbrella opening and the dog barking sound. Means and standard deviations are shown. The numbers above the bars are the rows in the data in which outliers occurred are recognized by SPSS.

During a full blink, vision is fully impaired, whereas, during a partial blink, vision is partially maintained. For the video analysis, Adobe Premiere Pro was used. The videos were cut into 10-second videos, put in greyscale, and exported in frames. The camera shot 50 frames per second, so about 500 frames were exported for each video. Then, the frames were analysed and the number of blinks was counted and marked down as full blinks (FB) or partial blinks (PB). A single observer scored all the blinks. Because eye blinks and head movements are often linked, it was also indicated whether each blink was associated with a head movement. Due to the quick movement of the heads of the chicks, and the frame rate of the camera, it could not be stated with certainty whether a blink occurred during a head movement. These occurrences (n=22) were left out of the data analysis. Statistical calculations were performed in R and SPSS.

## 3 Results

### 3.1 Blinking behaviour

**Figure 4:**
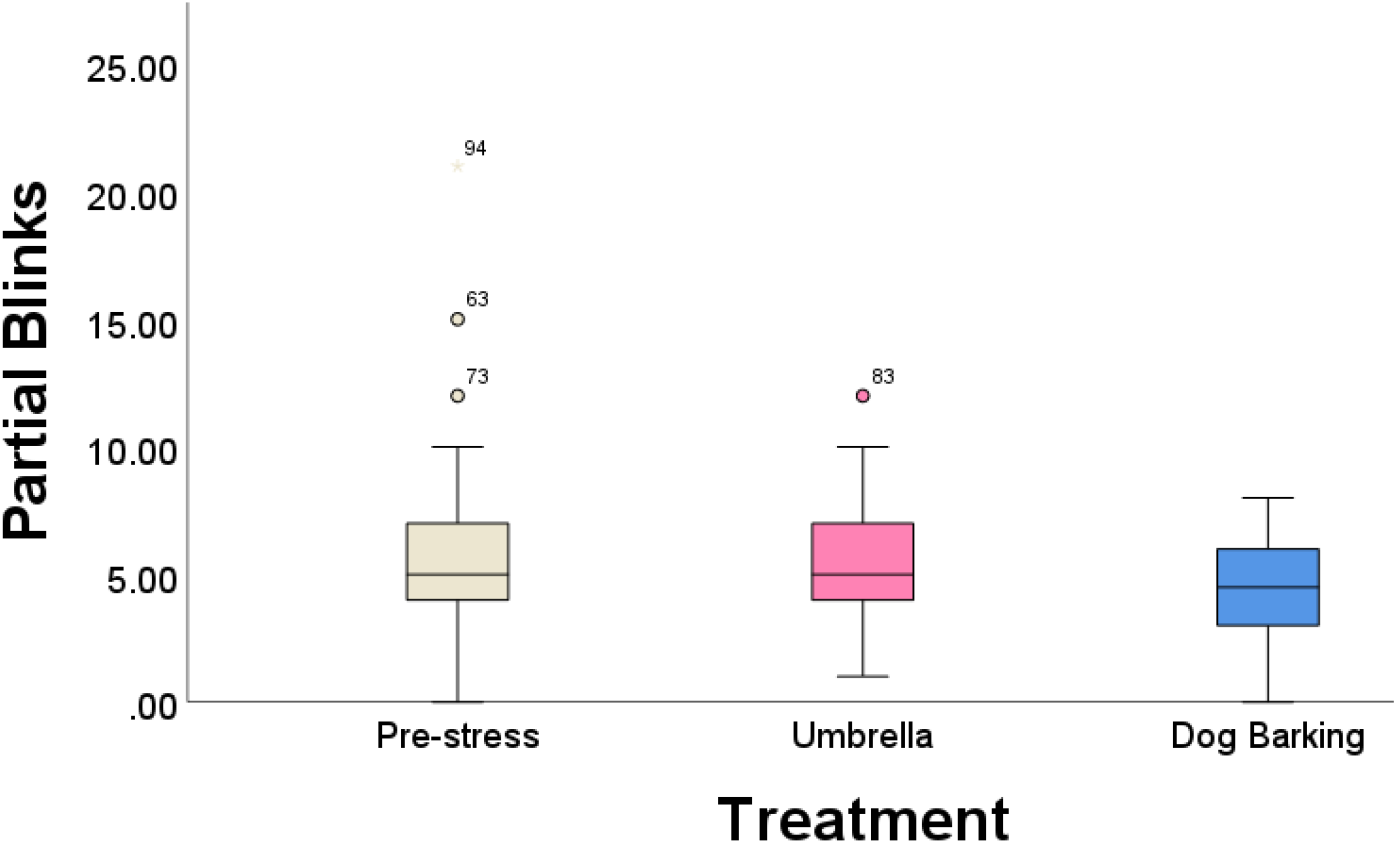
The number of partial blinks during the different treatments (Umbrella and Dog Barking). The average of partial blinks did not change significantly during pre-stress and stress treatments (p=0.065). Means and standard deviations are shown.

### 3.2 Thermal imaging

#### 3.2.1 Temperature and stress level

#### 3.2.2 Temperature and age

## 4 Discussion

This study aimed to investigate whether a negative affective state could be measured by assessing changes in blinking behaviour and temperature changes in the head region in response to stress treatments.

### 4.1. Blinking behaviour

Chickens possess two eyelids and a nictitating membrane on each eye. A nictitating membrane is a transparent third eyelid that is drawn horizontally across the eye. It can moisten the eye while maintaining vision (43). It keeps the cornea clean and can provide mechanical protection for the eye (44). Chickens do not typically blink with their upper and lower eyelid, but only with their nictitating membrane. The term “full blink” is therefore not necessarily used to refer to a blink, but is used as a term for a full closure of the eye. We looked at the eye closure as during this period they are fully non-vigilant. When a bird blink using only its nictitating membrane, it essentially still has partial vision. When a bird is stressed, thus needs full vigilance, it was expected blinking would still occur, as it is important for cleaning the eye and maintaining vision. However, complete eye closure fully impairs vision which is why it was decided to look at the frequency of these full blinks during stress as well.

The obtained results demonstrate that chicks do not significantly blink less partial blinks when they are stressed, but do display less full blinks when they are exposed to a stressor. This means the alertness increases during stress treatment. During the stress treatments with the visual stressor, the umbrella, the birds were respectively 9, 11, and 15 days old. In this period, the display of full blinks was inhibited. The stress treatments with the dog barking noise occurred when the birds were 17 and 19 days old. During this treatment, both a slight reduction in partial blinks and a reduction in full blinks were observed. The exposure of the chicks to the dog barking sound was an acute noise treatment. A study on the effect of noise on chicks investigated the neuroendocrine interaction during noise stress (45). Both the effects of acute noises and chronic noise were tested. Several physiological changes were measured, as well as behavioural changes related to water and feed intake. Loud noises of 80 and 100 dB caused more response during the acute noise treatment. For chronic stress treatment, the noises of 60 dB also evoked a response on water and feed intake. The present study did not measure the loudness of the dog barking sound used. It could be interesting in future studies to measure whether the loudness of the noises played affects the blinking behaviour in chicks.

Differences in the reaction to auditory and visual stress could be explained by the mechanisms of how chickens process information and how they react to stress. Chickens have a lateralized brain, which means information is processed differently in each brain hemisphere. The right hemisphere is more involved with fear, stress, predator detection, global spatial attention, and small differences in stimuli and the left hemisphere is more involved with the discrimination of objects from distracting stimuli, detection of large differences between stimuli, and proximal spatial attention (46). In this experiment, where the birds were exposed to auditory and visual stimuli that evoked a stress response, both hemispheres played an important role in processing information. Based on the number of blinks, it could not be detected whether one stress treatment gave a larger stress response than the other stress treatment, but the chicks did seem to react differently to the stressors in terms of full body behaviour. Behavioural changes in the whole body of the chicks could be observed in response to the stress treatments. These responses, such as increased wing flapping and increased vocalisations during the Umbrella treatment and standing response and reduced preening behaviour during both the Dog Barking treatment and the Umbrella treatment were not quantified during this study but could be interesting to explore in further research. Other studies identify behavioural stress responses in chicks as behavioural characteristics that differ from their normal performance. This could be aggressive behaviour, such as feather pecking or changes in feeding activity (5).

In addition, Yorzinski (2016) and Beauchamp (2017) found research that many blinks occur during gaze shifts, for example, when eating (35, 36). A study on the relationship between gaze shifts and blinking rate in peacocks found that the animals timed their blinks during large gaze shifts to concur with periods in which vision was already impaired. Blinking was inhibited mostly when they had to perform many gaze shifts, increasing their alertness (35). A study where the blinking rate was measured during feeding found that all blinks occurred when the birds made up and down movements while feeding. The blinks also occurred during gaze shifts, when the birds moved their heads from side to side to monitor their environment (36). The chickens blinked less often when they were observing their environment than when feeding. During feeding, the blinks were shorter than while monitoring. Despite the short blinks, the birds were still blind for nearly half the time when feeding.

As blinks are highly correlated with head movements and gaze shifts, it was important to quantify the number of blinks that did not occur during head movements. Out of the total number of blinks (PB and FB combined), 39.9% occurred without a gaze shift or head movement. The results (Figure 5) show a higher number of blinks without gaze shifts during prestress than during both treatments. They also show a higher amount of blinking without gaze shifts during the Dog Barking treatment than during the Umbrella treatment. The results could indicate that blinking rate decreases in stressful situations, independently from the head movements or gaze shifts. The duration of the blinks was not measured. It could be interesting to assess whether the partial blinks were shorter during stress than during prestress.

**Figure 5:**
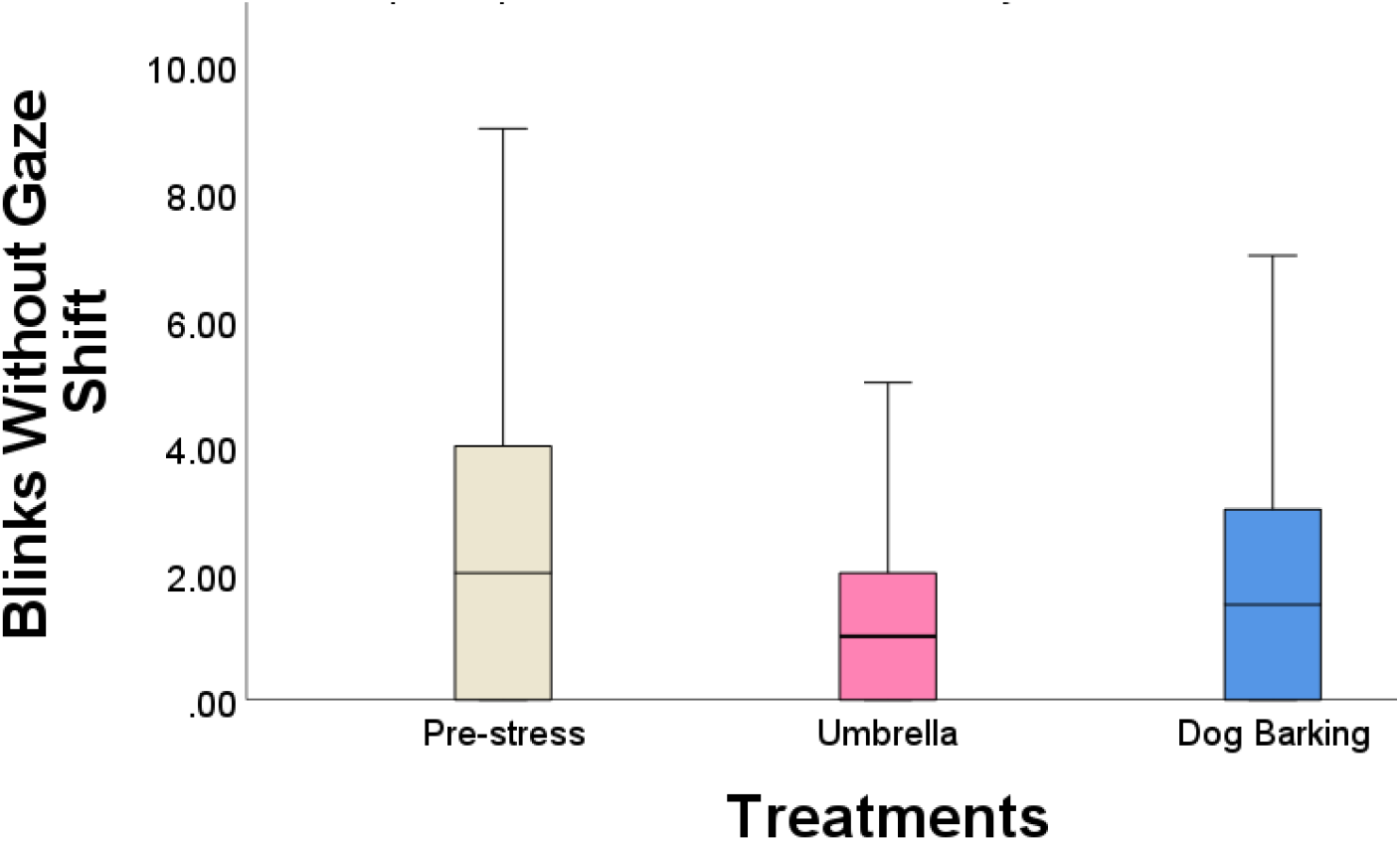
The number of blinks that occurred without a gaze shift during the different treatments (Pre-stress, Umbrella, and Dog Barking). The average of these blinks changed significantly between the three treatments (p=0.011). The average number of blinks changes significantly between pre-stress and the Umbrella treatment (p=0.04). There was also a significant change in the number of blinks between the Umbrella treatment and the Dog Barking treatment (p=0.037). No significant difference was found between pre-stress and the Dog Barking treatment (p=0.788). Means and standard deviations are shown.

**Figure 6:**
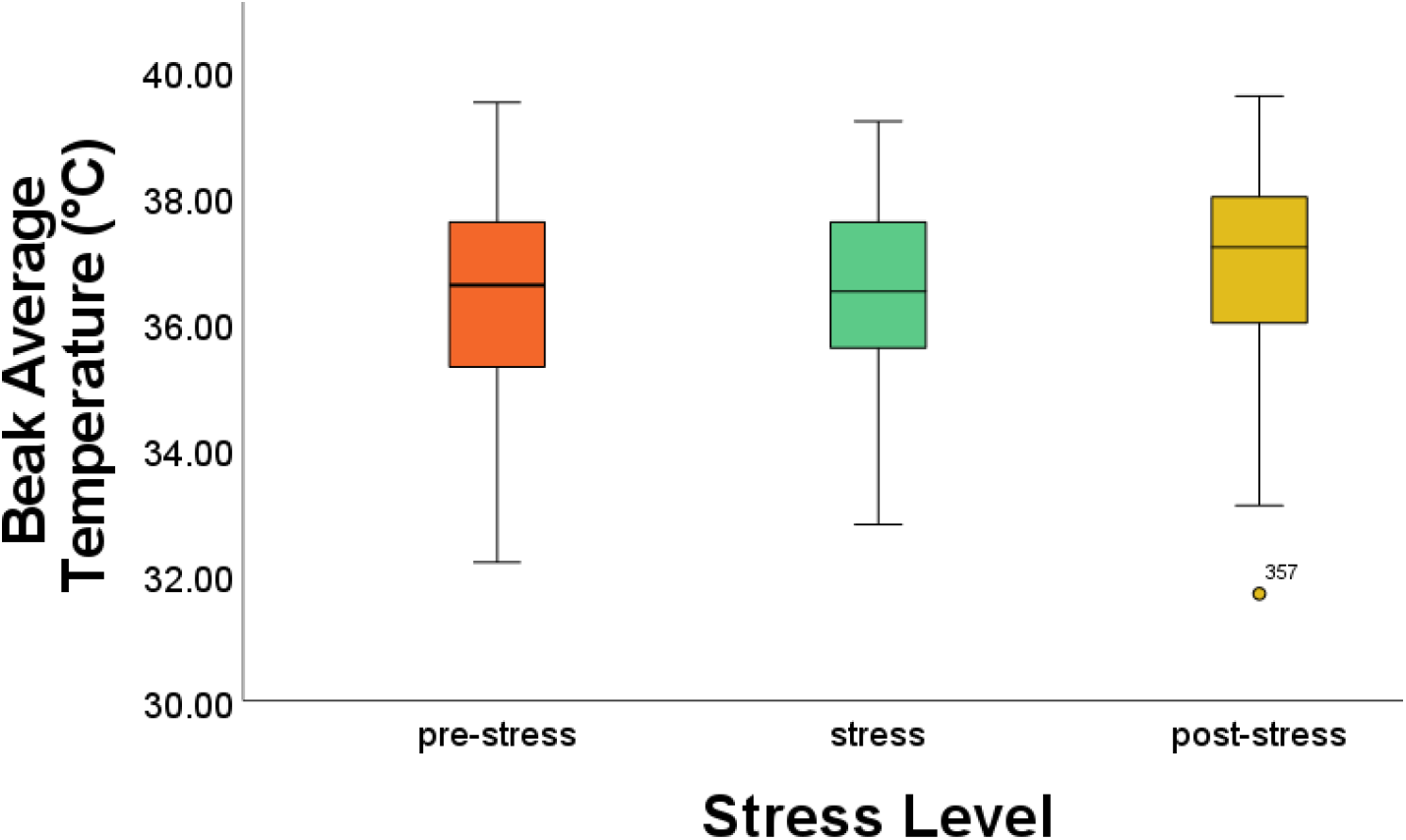
Boxplot of the mean beak temperature during the three levels of stress. There is a significant difference of 0.5 °C between pre-stress and post-stress (p=0.015). There is also a significant difference of 0.5 °C between stress and post-stress (p=0.024). No significant temperature difference was found during pre-stress and stress (p=0.871), adjusted for multiple comparisons. Means and standard deviations are shown.

**Figure 7:**
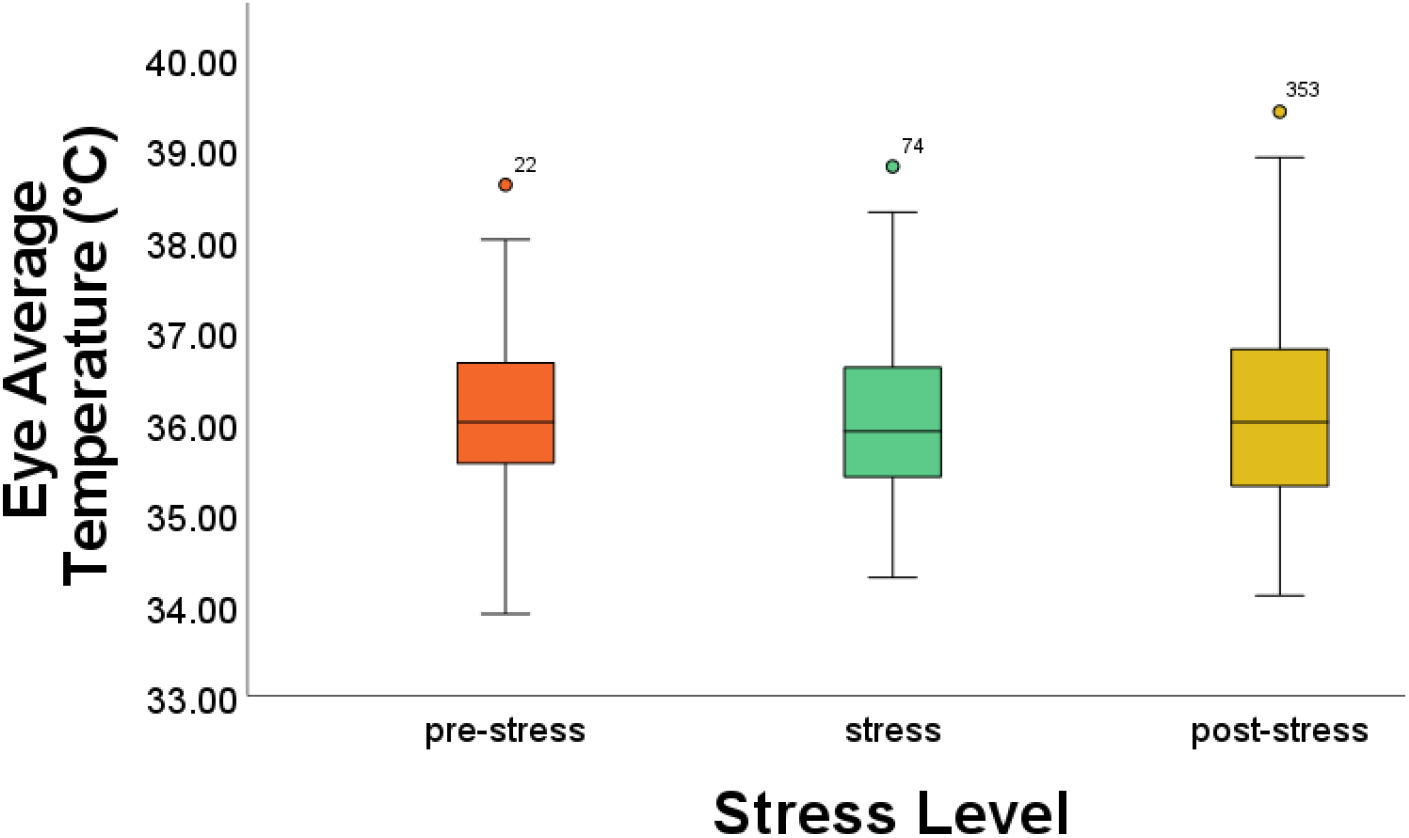
Boxplot of the mean eye temperature during the three levels of stress. For eye average temperature, no significant differences were found between pre-stress and stress (p=0.397), between stress and post-stress (p=0.396), and between pre-stress and post-stress (p=0.996). Means and standard deviations are shown.

**Figure 8:**
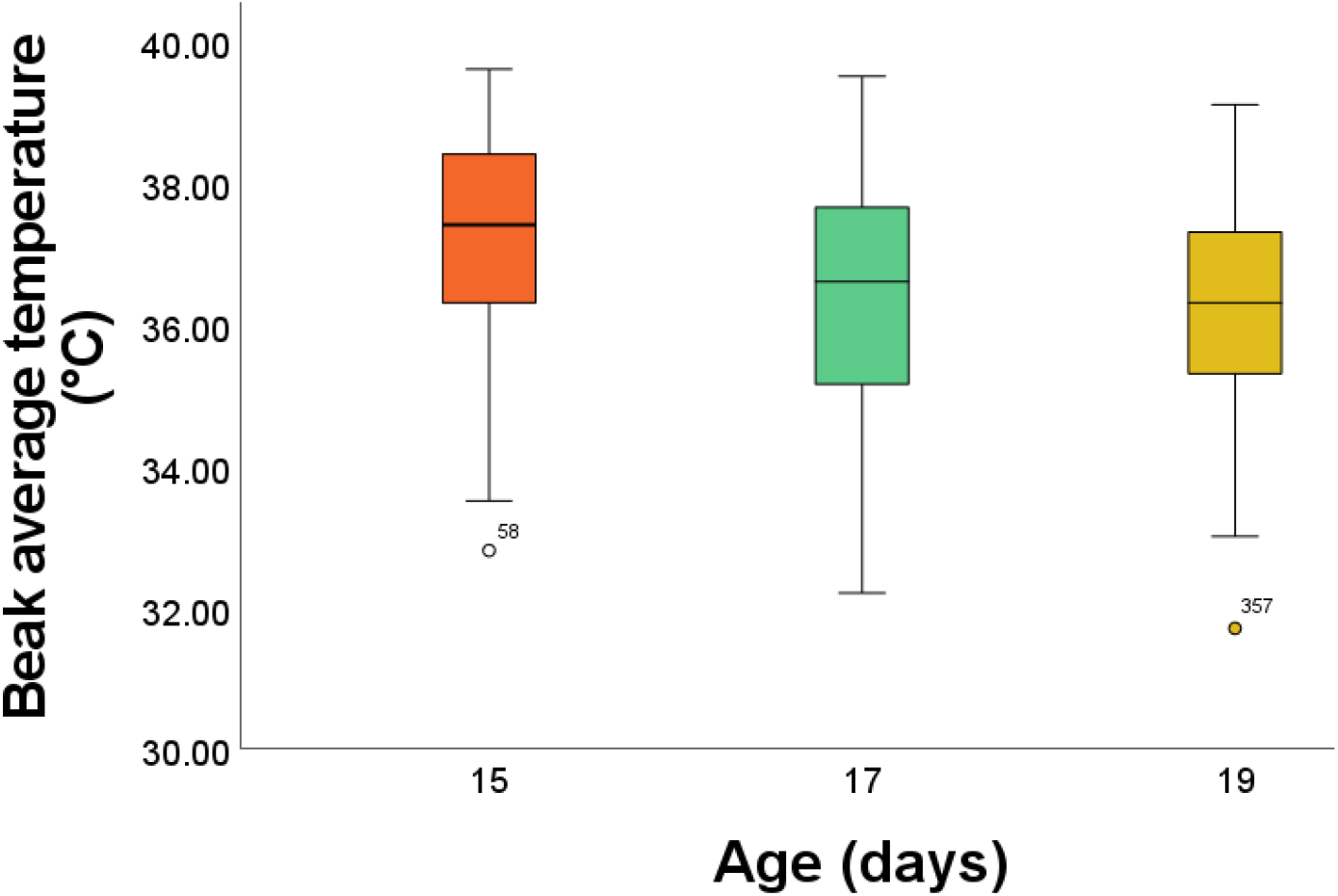
Boxplot of the beak average temperature during the different ages of the chicks. For beak average temperature, there is a significant mean difference of 0.78 °C between the 15- and 17-day old chicks (p=0.000) and a significant mean difference of 0.99 °C between the 15- and 19-day old chicks (p=0.000). No significant mean difference was found between the 17- and 19-day old chicks (p=0.289). Means and standard deviations are shown.

**Figure 9:**
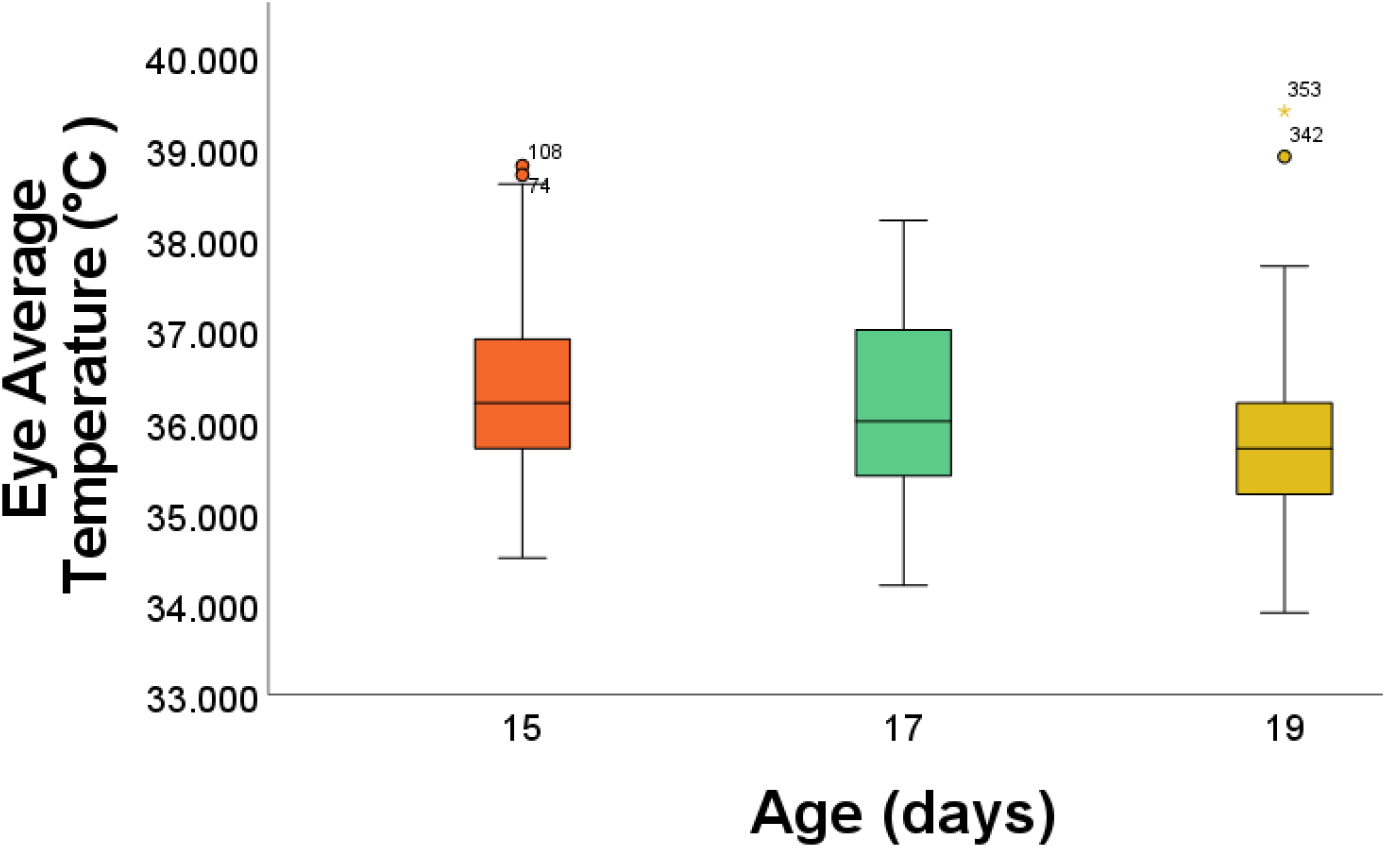
Boxplot of the eye average temperature during the different ages of the chicks. For eye average temperature, there is a significant mean difference of 0.60°C between the 15- and 19-day old chicks (p=0.000) and of 0.40 °C between the 17- and 19-day old chicks (p=0.002). There is no difference in average eye temperature between the 15- and 17-day old chicks (p=0.102). Means and standard deviations are shown.

As the chicks received the same treatments on multiple days (Umbrella treatment on days 9, 11, and 15 and Dog Barking treatment on days 17 and 19), there is a possibility that habituation may have taken place for these stress treatments. The results from the statistical analysis showed there was a significant difference in the number of blinks (Partial Blinks and Full Blinks combined) between days 9 and 19 (p=0.024), between days 11 and 17 (p=0.011), and between days 11 and 19 (p=0.002). However, the chicks were given different treatments on these days, which means effects cannot be attributed to habituation, but the different stress treatments. No significant differences were found in the number of blinks between the days on which the same stress treatment occurred. The chicks would have to be assessed for a longer period to observe whether habituation occurs.

### 4.2. Thermal imaging

This study assessed the changes in temperature in the head regions in response to a stressor. Chickens are homeotherm animals, meaning they keep a high and constant body temperature (47). However, when an animal experiences a certain affective state, such as fear or happiness, the body displays a physiological change (48).

The eye temperature showed a slight reduction in response to a stressor, though not significantly. This reduction in eye temperature as a response to a stressor was found in other studies where emotional contagion was measured within domestic chick broods (22), and in mother hens and chicks that were exposed to a stressor (24). A reduced eye temperature is an indicator of stress-induced hyperthermia (SIH) (22). Stress-induced hyperthermia, or emotional fever, is an acute stress response mediated by the activation of the hypothalamic-pituitary-adrenal axis (HPA-axis) in combination with sympathetic-adrenal medullary (SAM). This causes a release of stress hormones, such as cortisol, corticosterone, and temperature changes (49, 50).

In addition to the changes in eye temperature, the average beak temperature also changed in response to the stress treatments. The beak of birds is made of keratin. It is a very versatile organ, playing a role in many functions, such as feeding, drinking, preening, manipulating, nesting, and fighting (51). It possesses many nerve endings, is highly vascularised, and plays a role in thermoregulation (51). In toucans, the beak acts as a highly efficient thermoregulatory device and can exchange heat up to 400% of the resting heat production (52). Therefore, changes in beak temperature can be an interesting way of measuring affective states.

Feathers of birds are good insulators, trapping heat between the skin and the feathers. Although this is a beneficial trait for birds in colder climates, it also makes losing heat in warmer climates more difficult and makes the animals more susceptible to heat stress (53). A rise in temperature in the core body, for example, in response to a stressor, means this heat also must be dissipated to the outer environment. When a bird needs to get rid of heat, the beak and legs play a very important role (54).

Few studies have looked at the beak region temperature as a measure of stress. The present study found that the beak temperature significantly increased during the post-stress period, one hour after stress was induced. It is possible that due to the rise in core temperature, more heat had to be dissipated, which was done through the beak. Sometimes, the chicks had just drunk water before the thermographic pictures were taken. The results indicated a lower temperature in the beak, and, thus, a reduced mean temperature.

The head temperature of the chicks remained constant during the three levels of stress. The feathers on the head region of the chicks may have had an influence. The skin needs to be bare, which is not the case with the chicks’ heads.

The age of the birds has a significant influence on the eye temperature, the beak temperature, and the head temperature. Although the thermal camera was calibrated every day of use, there is a possibility environmental temperature played a role in the reduction of temperature in these three areas. As the birds got older, the environmental temperature reduced. At the start of the measurements, when the birds were nine days old, the environmental temperature had a maximum of 31.2°C. At the end of the measurement period, when the birds were 19 days old, the maximum environmental temperature was 27.9 °C. However, it is also possible the birds became less stressed, causing a smaller increase in core temperature, as they got older and adapted to the treatment.

In most studies involving thermographic imaging, the animals are photographed at a set distance, for example, one meter (55,29,56). As part of the study design, the chicks were always able to walk freely inside the cages and were not restrained. Thus, the thermographic pictures were taken at various distances and locations for each bird. This may have influenced the temperature of the chicks.

To validate the affective states measured in the laying hen chicks, studies on cortisol in feather and-or blood-based measures are needed. Acute and chronic stress can be assessed by measuring cortisol and corticosterone in, for example, feathers or blood (57,58). The advantage of blood measures is that hormonal changes in blood can happen quickly in response to a stressor, making it an effective way of measuring negative affective states (59). In feathers, corticosterone can also be measured but the levels do not change instantaneously, making it more difficult to assess affective states (59).

## 5 Conclusions

During this study, it was investigated whether affective states namely fear as an emotion could be assessed in laying hen chicks with the help of video and thermal imaging. The results showed that chicks display more full blinks before a stressor and that their blinking without gaze shifts decreased during a stressor. The reduction in the full blinking rate could indicate a negative affective state. The beak temperature changed significantly during the three different stress levels, in particular during post-stress. Stress-induced temperature changes in the beak and eye region of the chicks were significantly correlated to increase as the birds grew older. Future studies are warranted to explore whether the environmental temperature could play a role in the variation of temperatures or whether the chicks adapted to the stressors, causing less fearful responses. This study presented a quantitative examination of both physiological and behavioural changes in response to a stressor, which could be indicative of a negative affective state. For future research, feather or blood-based cortisol and corticosterone measurements are needed for validation of the negative affective states. With the advancements in AI enabled technology and sensors, blinking behaviour and the facial surface temperature can serve as new indicators for poultry welfare.

## Supporting information

Supplementary Videos and Thermal Videos Data

Supplementary Graphs Information

## 6 Supplementary Information

Sample video data showing the laying hens full blink head movement (SV1), full blink no head movement (SV2), full blink with head movement (SV3), head movement no blink (SV4), partial blink no head movement (SV5), partial blink with head movement (SV6) are available as supplementary files. Sample thermal video of laying hen chicks before stress (SV7 and SV8) and after stress by umbrella opening (SV9) are available under supplementary information. The results showing the correlation between the age of laying hen chicks to the total blinks and the stress type applied on the laying hen chicks to the head movement without blinks are available as figures S1 and S2 respectively. The results showing the correlation between the laying hen chick’s head average temperature to stress level and age of the chicks can be found as Supplementary graphs Figure S3 and S4 respectively.

## 8 Author Contributions

Conceptualization: SN; Methodology: SN, NP, HvH; Formal Analysis: NP, HvH; Investigation: NP, SN; Resources: SN; Writing – Original draft preparation: NP; Writing - review and editing: SN, JY, IM; Supervision, SN; Project administration: SN. Funding acquisition: SN.

## 10 Acknowledgments

The authors thank Bert Beukers from CARUS, animal experimental facility of the Wageningen University & Research for help with setting up animal care facilities and taking care of birds. The authors also thanks Stadsboerderij Wageningen (Petting Zoo) at the City of Wageningen, Netherlands for allowing us to test the thermal camera and video imaging equipment at their facilities.

## References

1. Giersberg MF, Poolen I, de Baere K, Gunnink H, van Hattum T, van Riel JW, et al. Comparative assessment of general behaviour and fear-related responses in hatchery-hatched and on-farm hatched broiler chickens. Appl Anim Behav Sci. (2020);232(105100):105100. doi: 10.1016/j.applanim.2020.105100

2. Edgar J, Held S, Paul E, Pettersson I, I’Anson Price R, Nicol C. Social buffering in a bird. Anim Behav. (2015);105:11–9. doi: 10.1016/j.anbehav.2015.04.007

3. Pittet F, Coignard M, Houdelier C, Richard-Yris M-A, Lumineau S. Effects of maternal experience on fearfulness and maternal behaviour in a precocial bird. Anim Behav. (2013);85(4):797–805. doi: 10.1016/j.anbehav.2013.01.026

4. Edgar J, Held S, Jones C, Troisi C. Influences of maternal care on chicken welfare. Animals (2016);6(1):2. doi: 10.3390/ani6010002

5. Hedlund L, Whittle R, Jensen P. Effects of commercial hatchery processing on short-and long-term stress responses in laying hens. Sci Rep. (2019);9(1):2367. doi: 10.1038/s41598-019-38817-y

6. Sih A, Bell AM, Johnson JC, Ziemba RE. Behavioral syndromes: An integrative overview. Q Rev Biol. (2004);79(3):241–77. doi: 10.1086/422893

7. Janczak AM, Riber AB. Review of rearing-related factors affecting the welfare of laying hens. Poult Sci. (2015);94(7):1454–69. doi: 10.3382/ps/pev123

8. Edgar J, Kelland I, Held S, Paul E, Nicol C. Effects of maternal vocalisations on the domestic chick stress response. Appl Anim Behav Sci. (2015);171:121–7. doi: 10.1016/j.applanim.2015.08.031

9. Wauters A-M, Richard-Yris M-A, Talec N. Maternal influences on feeding and general activity in domestic chicks. Ethology (2002);108(6):529–40. doi: 10.1046/j.1439-0310.2002.00793.x

10. Carew SN, Yusuf ND, Ejembi EP, Tuleun CD. Brooding and Rearing of Broiler Chicken on Free Range Using Local Chicken Hens. PAT. (2015);11(2):246–52.

11. Garnham L, Løvlie H. Sophisticated fowl: The complex behaviour and cognitive skills of chickens and red junglefowl. Behav Sci. (2018);8(1):13. doi: 10.3390/bs8010013

12. Marino L. Thinking chickens: a review of cognition, emotion, and behavior in the domestic chicken. Anim Cogn. (2017);20(2):127–47. doi: 10.1007/s10071-016-1064-4

13. Horback KM. The emotional lives of animals. In: The Routledge Handbook of Animal Ethics. Routledge; (2019);55–70.

14. Emery NJ, Clayton NS. Do birds have the capacity for fun? Curr Biol. (2015);25(1):R16–20. doi: 10.1016/j.cub.2014.09.020

15. Travain T, Valsecchi P. Infrared thermography in the study of animals’ emotional responses: A critical review. Animals (2021);11(9):2510. doi: 10.3390/ani11092510

16. Bertin A, Cornilleau F, Lemarchand J, Boissy A, Leterrier C, Nowak R, et al. Are there facial indicators of positive emotions in birds? A first exploration in Japanese quail. Behav Processes (2018);157:470–3. doi: 10.1016/j.beproc.2018.06.015

17. Anderson MG, Campbell AM, Crump A, Arnott G, Jacobs L. Environmental complexity positively impacts affective states of broiler chickens. Sci Rep. (2021);11(1):16966. doi: 10.1038/s41598-021-95280-4

18. Anderson MG, Campbell AM, Crump A, Arnott G, Newberry RC, Jacobs L. Effect of environmental complexity and stocking density on fear and anxiety in broiler chickens. Animals (2021);11(8):2383. doi:10.3390/ani11082383

19. Campbell DLM, Taylor PS, Hernandez CE, Stewart M, Belson S, Lee C. An attention bias test to assess anxiety states in laying hens. Peer J. (2019);7(e7303):e7303. doi: 10.7717/peerj.7303

20. de Waal FBM. Putting the altruism back into altruism: the evolution of empathy. Annu Rev Psychol. (2008);59(1):279–300. doi: 10.1146/annurev.psych.59.103006.093625

21. Nakamori T, Maekawa F, Sato K, Tanaka K, Ohki-Hamazaki H. Neural basis of imprinting behavior in chicks. Dev Growth Differ. (2013);55(1):198–206. doi: 10.1111/dgd.12028

22. Edgar JL, Nicol CJ. Socially-mediated arousal and contagion within domestic chick broods. Sci Rep. (2018);8(1). doi: 10.1038/s41598-018-28923-8

23. Tabh JKR, Burness G, Wearing OH, Tattersall GJ, Mastromonaco GF. Infrared thermography as a technique to measure physiological stress in birds: Body region and image angle matter. Physiol Rep. (2021);9(11):e14865. doi: 10.14814/phy2.14865

24. Edgar JL, Lowe JC, Paul ES, Nicol CJ. Avian maternal response to chick distress. Proc Biol Sci. (2011);278(1721):3129–34. doi:10.1098/rspb.2010.2701

25. Herborn KA, Graves JL, Jerem P, Evans NP, Nager R, McCafferty DJ, et al. Skin temperature reveals the intensity of acute stress. Physiol Behav. (2015);152(Pt A):225–30. doi: 10.1016/j.physbeh.2015.09.032

26. Jerem P, Herborn K, McCafferty D, McKeegan D, Nager R. Thermal imaging to study stress non-invasively in unrestrained birds. J Vis Exp. (2015);(105):e53184. doi: 10.3791/53184

27. Edgar JL, Paul ES, Nicol CJ. Protective mother hens: cognitive influences on the avian maternal response. Anim Behav. (2013);86(2):223–9. doi: 10.1016/j.anbehav.2013.05.004

28. Moe RO, Bohlin J, Flø A, Vasdal G, Stubsjøen SM. Hot chicks, cold feet. Physiol Behav. (2017);179:42–8. doi: 10.1016/j.physbeh.2017.05.025

29. Moe RO, Stubsjøen SM, Bohlin J, Flø A, Bakken M. Peripheral temperature drop in response to anticipation and consumption of a signaled palatable reward in laying hens (Gallus domesticus). Physiol Behav. (2012);106(4):527–33. doi: 10.1016/j.physbeh.2012.03.032

30. Marx G, Leppelt J, Ellendorff F. Vocalisation in chicks (Gallus gallus dom.) during stepwise social isolation. Appl Anim Behav Sci. (2001);75(1):61–74. doi: 10.1016/s0168-1591(01)00180-0

31. Mott RO, Hawthorne SJ, McBride SD. Blink rate as a measure of stress and attention in the domestic horse (Equus caballus). Sci Rep. (2020);10(1):21409. doi: 10.1038/s41598-020-78386-z

32. Marzluff JM, Miyaoka R, Minoshima S, Cross DJ. Brain imaging reveals neuronal circuitry underlying the crow’s perception of human faces. Proc Natl Acad Sci USA. (2012);109(39):15912–7. doi: 10.1073/pnas.1206109109

33. Gähwiler S, Bremhorst A, Tóth K, Riemer S. Fear expressions of dogs during New Year fireworks: a video analysis. Sci Rep. (2020);10(1):16035. doi: 10.1038/s41598-020-72841-7

34. Yorzinski JL. A songbird inhibits blinking behaviour in flight. Biol Lett. (2020);16(12):20200786. doi: 10.1098/rsbl.2020.0786

35. Yorzinski JL. Eye blinking in an avian species is associated with gaze shifts. Sci Rep. (2016);6(1). doi: 10.1038/srep32471

36. Beauchamp G. Half-blind to the risk of predation. Front Ecol Evol. (2017);5. doi: 10.3389/fevo.2017.00131

37. Prescott NB, Jarvis JR, Wathes CM. Vision in the laying hen. In: Welfare of the laying hen Papers from the 27th Poultry Science Symposium of the World’s Poultry Science Association (UK Branch), Bristol, UK, July 2003. Wallingford: CABI; 2004. p. 155–64.

38. Boissy A, Manteuffel G, Jensen MB, Moe RO, Spruijt B, Keeling LJ, et al. Assessment of positive emotions in animals to improve their welfare. Physiol Behav. (2007);92(3):375–97. doi: 10.1016/j.physbeh.2007.02.003

39. Duncan IJH, Filshie JH. The use of radiotelemetry devices to measure temperature and heart rate in domestic fowl. In: A handbook on biotelemetry and radio tracking, Oxford, Pergamon Press, Eds CJ Amlaner and DW MacDonald; 1980. p. 579–588.

40. Speakman J. Infrared thermography□: principles and applications. Zoology - analysis of complex systems. (2013) [cited 2022 Jan 5]; Available from: https://www.academia.edu/4686615/Infrared_thermography_principles_and_applications

41. Edgar JL, Nicol CJ, Pugh CA, Paul ES. Surface temperature changes in response to handling in domestic chickens. Physiol Behav. (2013);119:195–200. doi: 10.1016/j.physbeh.2013.06.020

42. Knoch, S., Whiteside, M. A., Madden, J. R., Rose, P. E., & Fawcett, T. W. (2021). Hot-headed peckers: thermographic changes during aggression among juvenile pheasants (Phasianus colchicus)

43. Heralgi M, Thallangady A, Venkatachalam K, Vokuda H. Persistent unilateral nictitating membrane in a 9-year-old girl: A rare case report. Indian J Ophthalmol. (2017);65(3):253–5. doi: 10.4103/ijo.IJO_436_15

44. Curio E. Wie Vögel ihr Auge schützen: Zur Arbeitsteilung von Oberlid, Unterlid und Nickhaut. J fur Ornithol. (2001);142(3):257–72. doi: 10.1007/bf01651365

45. Ray PP, Chatterjee T, Roy S, Rakshit S, Bhowmik M, Guha J, et al. Noise induces hypothyroidism and gonadal dysfunction via stimulation of pineal–adrenal axis in chicks. Proc Zool Soc. (2018);71(1):30–47. doi: 10.1007/s12595-016-0180-0

46. Rogers, L. J., & Kaplan, G. (2019). Does functional lateralization in birds have any implications for their welfare?. Symmetry, 11(8), 1043.

47. Jerem P, Jenni-Eiermann S, Herborn K, McKeegan D, McCafferty DJ, Nager RG. Eye region surface temperature reflects both energy reserves and circulating glucocorticoids in a wild bird. Sci Rep. (2018);8(1):1907. doi: 10.1038/s41598-018-20240-4

48. Kremer L, Klein Holkenborg SEJ, Reimert I, Bolhuis JE, Webb LE. The nuts and bolts of animal emotion. Neurosci Biobehav Rev. (2020);113:273–86. doi: 10.1016/j.neubiorev.2020.01.028

49. Charmandari E, Tsigos C, Chrousos G. Endocrinology of the stress response. Annu Rev Physiol. (2005);67(1):259–84. doi:10.1146/annurev.physiol.67.040403.120816

50. Ouyang JQ, Macaballug P, Chen H, Hodach K, Tang S, Francis JS. Infrared thermography is an effective, noninvasive measure of HPA activation. Stress (2021);24(5):584–9. doi: 10.1080/10253890.2020.1868431

51. Iqbal A, Moss AF. Review: Key tweaks to the chicken’s beak: the versatile use of the beak by avian species and potential approaches for improvements in poultry production. Animal (2021);15(2):100119. doi: 10.1016/j.animal.2020.100119

52. Tattersall GJ, Andrade DV, Abe AS. Heat exchange from the toucan bill reveals a controllable vascular thermal radiator. Science (2009);325(5939):468–70. doi: 10.1126/science.1175553

53. Mota-Rojas D, Titto CG, de Mira Geraldo A, Martínez-Burnes J, Gómez J, Hernández-Ávalos I, et al. Efficacy and function of feathers, hair, and glabrous skin in the thermoregulation strategies of domestic animals. Animals (2021);11(12):3472. doi: 10.3390/ani11123472

54. Fan L, Cai T, Xiong Y, Song G, Lei F. Bergmann’s rule and Allen’s rule in two passerine birds in China. Avian Res. (2019);10(1). doi: 10.1186/s40657-019-0172-7

55. Bartolomé E, Sánchez MJ, Molina A, Schaefer AL, Cervantes I, Valera M. Using eye temperature and heart rate for stress assessment in young horses competing in jumping competitions and its possible influence on sport performance. Animals (2013);7(12):2044–53. doi: 10.1017/S1751731113001626

56. Narayan E, Perakis A, Meikle W. Using thermal imaging to monitor body temperature of koalas (Phascolarctos cinereus) in A Zoo setting. Animals (2019);9(12):1094. doi: 10.3390/ani9121094

57. Freeman NE, Newman AEM. Quantifying corticosterone in feathers: validations for an emerging technique. Conserv Physiol. (2018);6(1):coy051. doi: 10.1093/conphys/coy051

58. Weimer SL, Wideman RF, Scanes CG, Mauromoustakos A, Christensen KD, Vizzier-Thaxton Y. An evaluation of methods for measuring stress in broiler chickens. Poult Sci. (2018);97(10):3381–9. doi: 10.3382/ps/pey204

59. Ataallahi M, Nejad JG, Song J-I, Kim J-S, Park K-H. Effects of feather processing methods on quantity of extracted corticosterone in broiler chickens. J Anim Sci Technol. (2020);62(6):884–92. doi: 10.5187/jast.2020.62.6.884

