## Supplementary Graphs Information for "Understanding Chicks’ Emotions: Are Eye Blinks & Facial Temperatures Reliable Indicators?"

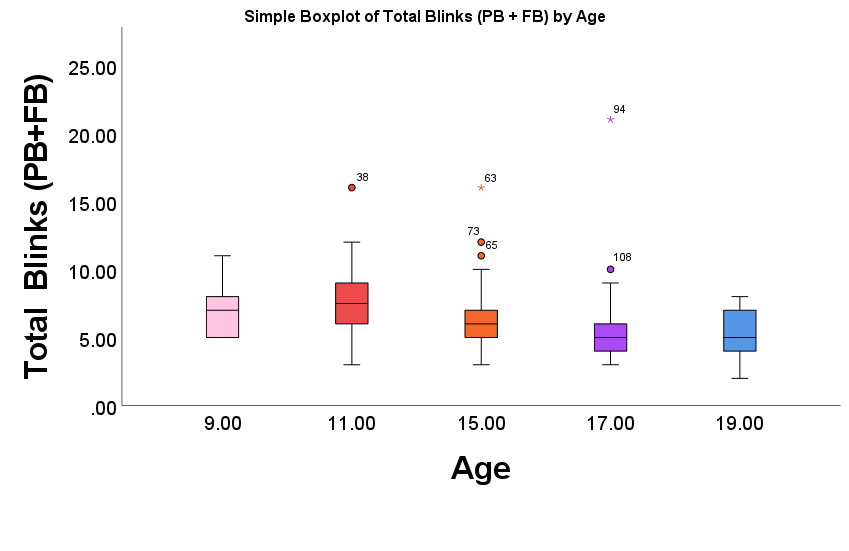

**Figure S1**: The total number of blinks in relation to age. There is a significant difference in the number of blinks (Partial Blinks and Full Blinks combined) between day 9 and 19 (p=0.024), between day 11 and 17 (p=0.011) and between day 11 and 19 (p=0.002). Means and standard deviation are shown.

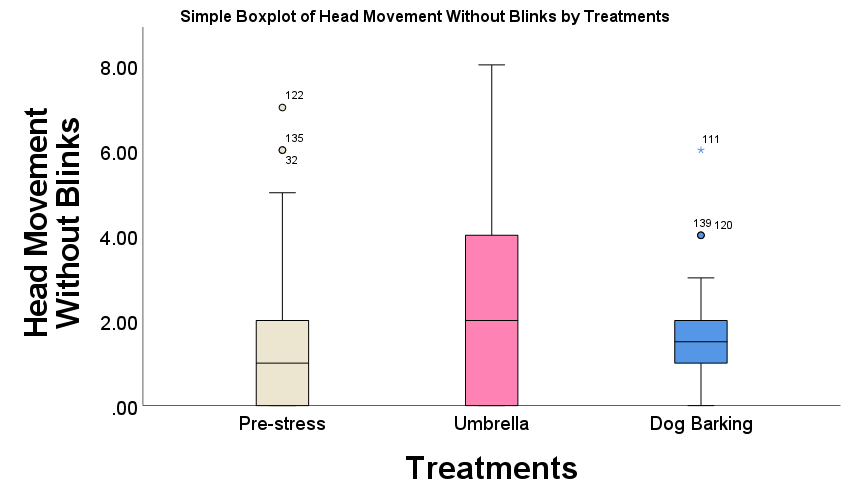

**Figure S2**: The number of head movements in relation to the three stress treatments. There is a significant difference in the number of head movements between Pre-stress and the Umbrella treatment (p=0.045). No significant difference was found between Pre-stress and the Dog Barking treatment (p=0.881) and between the Umbrella treatment and the Dog Barking treatment (p=0.082). Means and standard deviation are shown.

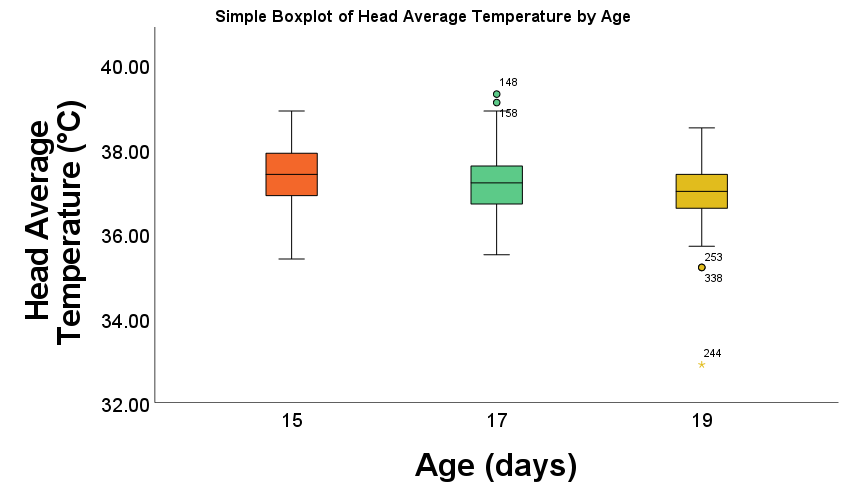

**Figure S3**: Boxplot of the mean head temperature during the three levels of stress. For head average temperature, no significant differences were found between pre-stress and stress (p=0.965), between stress and post-stress (p=0.881) and between pre-stress and post-stress (p=0.915). Means and standard deviations are shown.

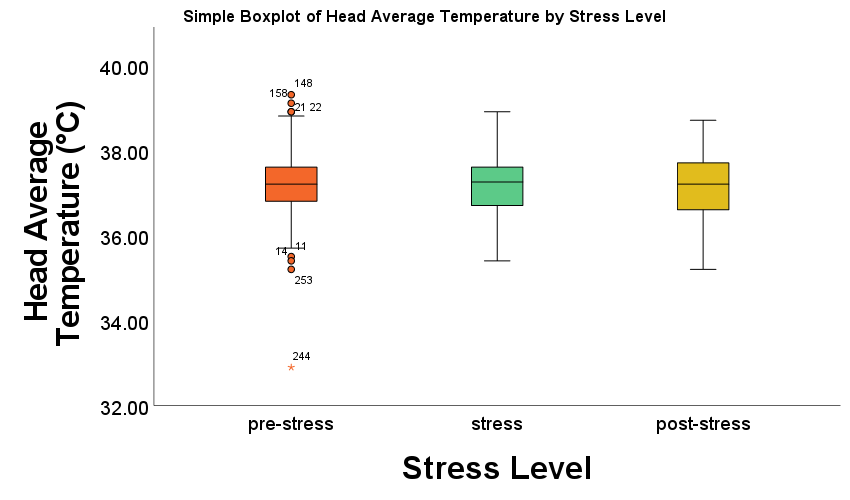

**Figure S4**: Boxplot of the head average temperature during the different ages of the chicks. For head average temperature, there is a significant mean difference of 0.39 °C between the 15- and 19-day old chicks (p=0.000) and of 0.21 °C between the 17- and 19-day old chicks (p=0.33). There is no difference in average eye temperature between the 15- and 17-day old chicks (p=0.063). Means and standard deviations are shown.
